# NMR-guided rational exploration of co-factors in boosting the *Pfu* DNA polymerase

**DOI:** 10.1101/2023.11.17.567503

**Authors:** Yihao Chen, Mingjun Zhu, Ruishen Ding, Xiaoling Zhao, Zhiqing Tao, Xu Zhang, Maili Liu, Lichun He

**Affiliations:** State Key Laboratory of Magnetic Resonance Spectroscopy and Imaging, National Center for Magnetic Resonance in Wuhan, Innovation Academy for Precision Measurement Science and Technology, Chinese Academy of Sciences, Wuhan 430071, China; University of Chinese Academy of Sciences, Beijing, 100049, China; Department of Reproductive Medicine, General Hospital of Central Theater Command of the People’s Liberation Army, Wuhan, Hubei 430061, China; Qinhe Life Science Ltd. Wuhan 430000, China; Optics Valley Laboratory, Hubei 430074, China

**Keywords:** *Pfu* DNA polymerase, co-factors, NMR Spectroscopy

## Abstract

With rapid developments of emerging technologies like synthetic biology, the demand for DNA polymerases with superior activities including higher thermostability and processivity has increased significantly. Thus, rational optimization of the performance of DNA polymerase is of great interest. Nuclear magnetic resonance spectroscopy (NMR) is a powerful technique used for studying protein structure and dynamics. It provides the atomic resolution information of enzymes under its functional solution environment to reveal the active sites (hot spots) of the enzyme, which could be further used for optimizing the performance of enzymes. In our previous work, we identified hot spot residues of *Pfu* DNA polymerase (*Pfu* pol). We aim to employ these binding hot spots to screen for co-factors of *Pfu* pol, particularly targeting those molecules exhibiting weak intermolecular interactions. To validate this concept, we first demonstrated the feasibility of utilizing hot spot residues as screening probes for auxiliary factors by employing the well-characterized Tween-20 as a model system. Employing these hot spots as probes, two new co-factors, the heat shock protein TkHSP20 from *Thermococcus Kodakaraensis* and the chemical chaperone L-arginine, are identified to interact with *Pfu* pol to boost its performance in amplifying long DNA fragments by enhancing the thermal stability and the processivity of the *Pfu* pol. This NMR-based approach requires no prior assignment information of target enzymes, guiding the rational exploration of novel co-factors for *Pfu* pol. Moreover, our approach is not dependent on structural data or bioinformatics. Therefore, it has significant potential for application in various enzymes to expedite the progress in enzyme engineering.

## 1. INTRODUCTION

PCR technology serves as the cornerstone for sequencing, molecular diagnostics, and synthetic biology [1-4]. Despite its apparent simplicity, certain challenges persist, particularly in the context of long-range PCR. One major problem in amplifying long DNA fragments is the long elongation time required at high temperatures [5], which will diminish the activity of DNA polymerases [6]. DNA polymerases must endure throughout the entire polymerase chain reaction to enable DNA amplification. Additionally, with the augmentation of DNA length, there is an increased probability of secondary structure formation and premature amplicon generation, potentially impeding the PCR process. DNA polymerase with higher processivity is required to address this issue. Several approaches have been developed to improve the amplification of lengthy DNA fragments, encompassing optimized buffer components, primer design, amplification protocols, and fusion domain engineering [7-10]. Of these approaches, the addition of cofactors directly aiding the DNA polymerase to facilitate the synthesis of long DNA fragments is critical.

Chaperones are highly conserved molecular machines that play a vital role in maintaining the functionality of the proteome under both normal and stressful conditions. They participate in various cellular processes, including the folding of newly synthesized proteins [11], the prevention of the aggregation of misfolded proteins [12,13], and the restoration of denatured proteins [14,15]. Chaperones essentially serve as guides for client proteins, reducing the likelihood of denaturation, thereby, prolonging the lifetime and activity of client proteins. Additionally, plenty of studies also suggest that small, low-molecular-weight compounds can be effective in inhibiting protein aggregation or enabling proteins to restore their functions [16,17]. These small molecules are referred to as chemical chaperones, and they are believed to non-specifically stabilize proteins, facilitating their proper folding. Among chemical chaperones, L-arginine (Arg) stands out as the most potent additive for suppressing protein aggregation induced by heat and reduced reagents, as well as aggregation during *in vitro* folding. Further to this, Bovine serum albumin (BSA), known for its chaperone-like properties [18,19], is widely employed in PCR systems [20-23], although its mechanism is not thoroughly clear. Thereby, we are intrigued by the potential functions of chaperones or other chaperone like additives in improving the performance of DNA polymerase. However, how to identify and prove the role of additives in aiding the activities of the DNA polymerase needs a rational detectable approach.

Nuclear magnetic resonance (NMR) spectroscopy is a powerful and versatile tool used for protein structure and dynamics analysis. It has the capability to decipher intricate structures and interactions of proteins in real-time, even within its native heterogeneous multiphase working environments, such as in cells and cell-like environments [24]. Thus, NMR holds immense promise in biomolecule research, offering profound insights into a wide spectrum of enzyme catalytic processes at the molecular level. This includes the identification of binding sites (hot spots) for specific substrates in the working solution and the exploration of changes of hot spots on both enzyme and its substrates. We based our approach on the hot spots identified in our previous work [25]. By targeting these hot spots, we identified the chemical chaperone, L-arginine, and the heat shock protein, HSP20 from *Thermococcus kodakaraensis* engaged with *Pfu* pol to bolster its thermal resilience, processivity and enhance its activity in amplifying lengthy DNA fragments using 2D [^13^C,^1^H]-HMQC. This approach guides the rational exploration of novel additives for *Pfu* pol and demonstrates significant potential for application in other enzymes, thereby expediting progress in enzyme engineering.

## 2. MATERIAL AND METHODS

### 2.1 Cloning, expression, and purification of proteins

The genes encoding *Pfu* DNA polymerase (*Pfu* pol) and *Pfu* DNA polymerase*-*Sso7d polymerase (*Pfu* pol-S) were codon-optimized, synthesized by Sangon Biotech Co., Ltd., and cloned into the pET16b vector (Novagen) with an N-terminal His-tag. Similarly, the *TkHSP20* gene was codon-optimized, synthesized by Sangon Biotech Co., Ltd., and inserted into a modified pET21a vector fused with a 10×His -SUMO tag using FasHifi Super DNA polymerase (Affinibody Lifescience Co., Ltd.). The respective plasmids were introduced into BL21 (DE3) competent cells (Novagen), which were cultured in LB broth or M9 medium supplemented with 100 mg/mL ampicillin. Protein expression was induced with 0.5 mM isopropyl-β-D-thiogalactopyranoside (IPTG) at an optical density (600 nm) of 0.6, followed by incubation at 37 °C for 6 hours. Uniformly [^13^C] -labeled protein was obtained by growing cells in M9 medium supplemented with [U-^13^C] glucose (3 g/L) [26]. Cells were harvested by centrifugation at 5000 × g for 25 minutes at 4 °C.

For *Pfu* pol and *Pfu* pol-S, the cell pellet was resuspended in 50 mL of Lysis Buffer A (pH 9.0, 50 mM Tris-HCl, 500 mM KCl, 0.1 mM PMSF, and 0.01 mg/mL DNase I). For SUMO-TkHSP20, the pellet was resuspended in 50 mL of Lysis Buffer B (pH 9.0, 50 mM Tris-HCl, 300 mM NaCl, 0.1 mM PMSF, and 0.01 mg/mL DNase I). Cell lysis was performed using a high-pressure homogenizer, followed by centrifugation at 27,000 × g for 30 minutes at 4 °C to separate the soluble fraction. For the purification of *Pfu* pol and *Pfu* pol*-S*, the supernatant was incubated at 70 °C for 30 minutes to precipitate host proteins from *E. coli*, followed by an additional centrifugation step at 27,000 × g for 30 minutes. The clarified supernatant containing the target protein was then loaded onto a Ni-NTA (nitrilotriacetic acid) affinity column (Qiagen) pre-equilibrated with the corresponding lysis buffer. The column was washed with 100 mL of Wash Buffer (each respective Lysis Buffer supplemented with 30 mM imidazole), and *Pfu* pol or *Pfu* pol*-S* was subsequently eluted with 25 mL of Elution Buffer A (pH 8.0, 50 mM Tris-HCl, 50 mM KCl, 300 mM imidazole). For SUMO-TkHSP20, the bound protein was eluted with 25 mL of Elution Buffer B (pH 9.0, 50 mM Tris-HCl, 300 mM NaCl, 300 mM imidazole). For further purification of *Pfu* pol and *Pfu* pol*-S*, the eluted fractions were applied to a 5 mL HiTrap Heparin column (GE Bioscience) pre-equilibrated with Heparin Buffer (pH 8.0, 50 mM Tris-HCl, 50 mM KCl). The proteins were eluted using a 0–500 mM KCl gradient in Heparin Buffer (pH 8.0, 50 mM Tris-HCl). The purified proteins were immediately buffer-exchanged into Storage Buffer (pH 7.4, 25 mM Tris-HCl, 50 mM KCl, 0.1 mM EDTA, 1 mM DTT, 50% glycerol) using a PD-10 column (GE Bioscience) and stored at -20 °C. For SUMO-TkHSP20, the eluted fractions were dialyzed against SEC Buffer A (pH 9.0, 50 mM Tris-HCl, 300 mM NaCl) supplemented with 1 mM DTT. SUMO-tag cleavage was performed overnight at 4 °C using SUMO protease. The untagged TkHSP20 was separated using a reverse Ni-NTA column, and the flow-through fractions containing TkHSP20 were collected. Further purification was carried out using a Superdex-200 size-exclusion chromatography column (GE Bioscience) equilibrated in SEC Buffer A. The purified TkHSP20 was concentrated and stored at -80 °C for future use.

### 2.2 NMR Titration Experiments

All NMR experiments involving *Pfu* pol were conducted in NMR buffer (20 mM sodium phosphate, 35 mM NaCl, 0.02% NaN3, 8% D2O, pH 7.4). Unless otherwise specified, all spectra were acquired at 298K using a Bruker Avance 600 MHz NMR spectrometer equipped with a cryogenically cooled probe. To investigate the interaction between *Pfu* pol and the selected additives, 2D [^13^C,^1^H] -HMQC spectra were recorded for 50 μM uniformly ^13^C-labeled *Pfu* pol in the presence of either 20 mM L-arginine or 0.05% Tween-20. The selection of these conditions was based on previously identified hotspot residues. L-arginine was prepared in phosphate buffer at pH 7.4 to prevent the introduction of additional salt ions. For all NMR spectra, 1024 complex points were recorded in the direct dimension, and 154 complex points were collected in the indirect dimension. Data processing included the application of a shifted sine bell function and polynomial baseline correction in all dimensions. NMR data were processed and analyzed using CCPNMR Analysis [27]. The combined chemical shift perturbation (CSP) (δ_comb_) for each residue was calculated using the following equation:

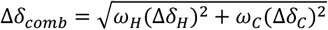

where *Δδ*_*H*_ and *Δδ*_*C*_ represent chemical shift changes (in ppm) in the ^1^H and ^13^C dimensions, respectively, while *ω*_*H*_ and *ω*_*C*_ are normalization factors (*ω*_*H*_ = 1.00, *ω*_*C*_ = 0.34 for aliphatic carbons or 0.07 for aromatic carbons). The binding affinity between *Pfu* pol and TkHSP20 was determined through NMR titration. Spectra of 50 μM ^13^C-labeled *Pfu* pol were acquired in the presence of increasing concentrations of TkHSP20 (0, 25, 50, 100, and 200 μM) at both 298K and 313K. The normalized CSP magnitude for residue aro3 of *Pfu* pol was plotted as a function of TkHSP20 concentration, and the dissociation constant (*K*_d_) was obtained by fitting the binding curve according to a previously established protocol [28].)

### 2.3 Fluorescence-based thermal shift assay

Thermal shift experiments were conducted using a QuantStudio 3 Real-Time PCR instrument (Thermo Fisher Scientific). The melting temperature was programmed to increase from 25 °C to 98 °C in 0.15 °C increments. Each reaction was carried out in a final volume of 20 μL containing 50 mM Tris-HCl (pH 8.8), 10 mM KCl, 10× SYPRO Orange (Sigma), and 10 µM of the respective protein. Fluorescence signals were recorded every second. To assess the effects of L-arginine and TkHSP20 on the stability of *Pfu* pol, samples were incubated at 95 °C for 120 minutes while continuously monitoring fluorescence signals. The recorded fluorescence intensities were plotted against time, and the first derivative of the fluorescence signal curve was calculated using GraphPad Prism 8.

### 2.4 Endpoint PCR

PCR reactions (20 μL) were prepared by mixing 2× buffer with the respective polymerase reaction buffer: *Pfu* pol reaction buffer (pH 8.8, 50 mM Tris-HCl, 10 mM KCl, 6 mM ammonium sulfate, 2 mM MgCl2, 0.05% Triton X-100, 0.001% BSA) or *Pfu* pol*-S* pol reaction buffer (pH 8.8, 50 mM Tris-HCl, 100 mM KCl, 2 mM MgCl2, 0.05% Triton X-100, 0.001% BSA). Each reaction contained 50 nM *Pfu* pol or *Pfu* pol*-S*, 4 ng plasmid DNA, 50 ng Alpaca cDNA, 100 ng mouse tail genomic DNA, or 30 ng λ DNA as the template, along with 2 μM of each primer (see Table S1), 0.2 mM dNTPs, and varying concentrations of additives as specified in the figure legends. PCR amplification was carried out using a C1000 Touch thermal cycler (Bio-Rad). The reaction was initiated with an initial denaturation at 95 °C for 3 minutes. For *Pfu* pol, amplification proceeded through 33 cycles of 95 °C for 30 seconds, 55 °C for 30 seconds, and 72 °C f or 45 seconds to 10 minutes, depending on the target DNA length (1 kb/min). For *Pfu* pol*-S*, the reaction consisted of 33 cycles of 95 °C for 15 seconds, 60 °C for 15 seconds, and 72 °C for 20 seconds to 5 minutes, with an extension rate of 2 kb/min. Following amplification, 4 μL of the PCR product was mixed with loading dye and subjected to electrophoresis on a 1% or 0.6% agarose-TAE gel containing 1× GenRed nucleic acid gel stain (Genview). Gels were visualized under ultraviolet light, and the band intensity of PCR products was quantified using ImageJ. The resulting intensity plots were generated using GraphPad Prism 8.

### 2.5 Thermal stability PCR assay

Apo-enzyme (*Pfu* pol, 50 nM) and holo-enzyme (*Pfu* pol, 50 nM, supplemented with 100 pg/μL TkHSP20) were incubated at 72 °C for varying durations in 2× *Pfu* pol reaction buffer (pH 8.8, 100 mM Tris-HCl, 20 mM KCl, 12 mM ammonium sulfate, 4 mM MgCl2). At each time point, a 25 μL aliquot was collected and mixed with 10 ng plasmid DNA template (target length: 10 kb), 0.2 μM of each primer, 0.2 mM dNTPs, and ddH2O to a final reaction volume of 50 μL. PCR amplification, gel electrophoresis, and data analysis were performed as described previously. ImageJ was used to quantify the intensity of PCR product bands, and GraphPad Prism 8 was utilized to generate intensity plots, allowing estimation of the half-life of *Pfu* pol in the presence and absence of 100 pg/μL TkHSP20.

### 2.6 Steady-state kinetic analyses

Steady-state kinetic data were collected and analyzed using an EvaGreen-based fluorometric polymerase activity assay [29]. Hairpin DNA (sequence provided in Table S1) was prepared at a concentration of 100 μM in water by heating at 98 °C for 5 minutes, followed by annealing on ice for 30 minutes. Varying concentrations of hairpin DNA (0.1–2 μM) were then used as templates and mixed with dNTPs and 2 nM *Pfu* pol in the presence or absence of 2 mM L-L-arginine. DNA synthesis was initiated by the addition of MgCl2 at 72 °C. The reaction buffer consisted of 25 mM Tris-HCl (pH 8.8), 10 mM KCl, 2.5 mM MgCl2, 200 μM dNTPs, and 0.05% Triton X-100. Fluorescence intensity was recorded over time, with background subtraction applied to each reaction to obtain the corrected fluorescence values. The initial reaction rate for each template concentration was determined by calculating the first derivative of the fluorescence curve. Kinetic parameters (*k*_*cat*_ and *K*_m_) were derived by fitting the Michaelis-Menten equation using GraphPad Prism 8.0. Reported values represent the mean of two biological replicates, each performed with three technical replicates.

## 3. RESULTS

### 3.1 Hot spots can be used for co-factors screening

In our previous experiments, we identified the active hot spots (ali1, aro1, aro2, aro3, and aro4) of *Pfu* pol by analyzing the extent of CSPs in the side-chain signals of hot spot residues in the bispecific nanobody and the specific hairpin substrate, based on NMR 2D ^1^H -^13^C HMQC spectra [25]. During the study, we observed that the previously reported cofactor Tween-20 also induced CSPs in the active hot spot residues to a certain extent (Fig. 1A). The addition of 0.01-0.1% Tween-20 into the *Pfu* pol reaction system could enhance its performance in amplifying different templates with varied length (Fig.1C, D, and E). Meanwhile, NMR data in Fig. 1B revealed the addition of 0.05% Tween-20 into the *Pfu* pol resulted in CSPs of the residues ali1, aro1, aro3, and aro4. These results indicated Tween-20 interacted with *Pfu* pol on the hot spots, strengthening the applicability of these hot spots as probes for identifying co-factors in boosting the *Pfu* pol.

**Figure 1.**
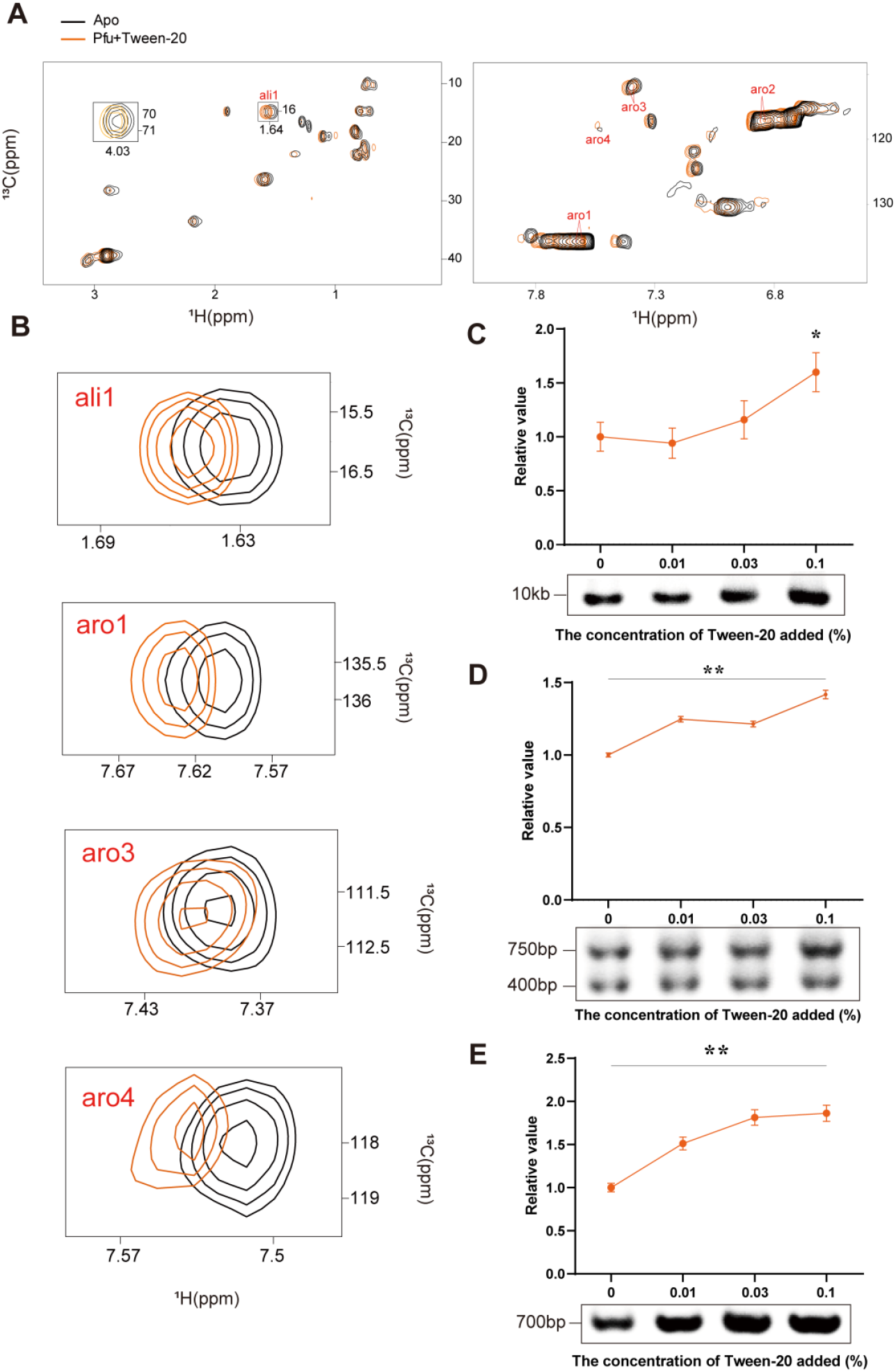
**(A)** 2D [^13^C,^1^H]-HMQC spectra of 50 μM [^13^C]-labeled *Pfu* pol acquired on the 600 MHz (^1^H frequency) spectrometer at 298 K in the absence (black) and presence (orange) of 0.05% Tween-20 (**A**). (**B**) Plots of chemical shift perturbations of ‘hot spots’ of 50 μM [^13^C]-labeled *Pfu* pol upon adding 0.05% Tween-20. (**C, D** and **E**) PCR amplification of different DNA template by the *Pfu* pol in the presence of different target length. PCR amplification of 10 kb plasmid DNA substrate (**C**), Alpaca cDNA substrate (**D** with target length 750 bp and 400 bp) and Mouse tail gDNA substrate (**E**, with target length 700 bp). Intensities of gel bands were quantified and plotted against the concentration of Tween-20, showing on top of the corresponding gel image.

### 3.2 Both L-arginine and TkHSP20 enhance the performance of *Pfu* pol in long-range PCR

We extended our study to explore novel additives for *Pfu* pol and investigated the effects of L-arginine, a chemical chaperone commonly used for protein stabilization and refolding [30,31]. Preliminary tests revealed that L-arginine (1–14 mM) had negligible impact on the amplification of short DNA fragments (Fig. 2A and S2A), and to the best of our knowledge, no prior reports have described its enhancement of PCR efficiency. However, significant chemical shift perturbations (CSPs) were observed at key residues ali1, aro1, aro2, and aro4 upon L-arginine addition (Fig. 2B and S1A), prompting further investigation. The application of L-arginine in long range PCR was then tested. Notably, in the amplification of longer DNA templates (10 kb), endpoint PCR assays demonstrated that L-arginine significantly improved *Pfu* pol yield in a concentration-dependent manner, with a ∼30% increase in final PCR product at 2 mM (Fig. 2C, D).

**Figure 2.**
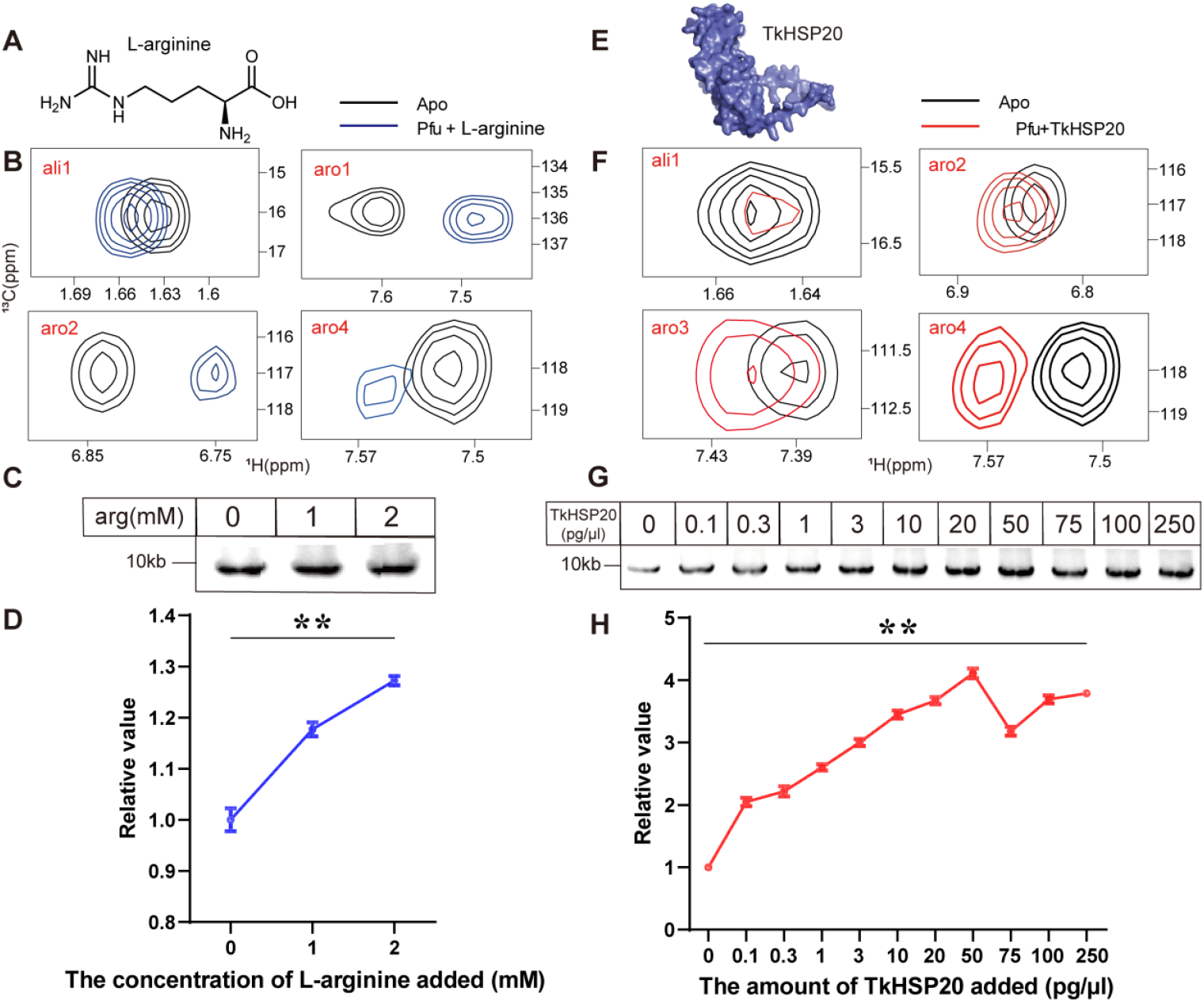
L-arginine and TkHSP20 enhances the performance of *Pfu* pol in the long-range PCR. (**A**) Chemical structures of L-arginine. (**B**) Plots of chemical shift perturbations of ‘hot spots’ of 50 μM [^13^C]-labeled *Pfu* pol upon adding 20 mM L-arginine. (**C**) PCR amplification of 10 kb plasmid DNA by the *Pfu* pol in the presence of different concentrations of L-arginine from 1 mM to 2 mM. PCR assays were performed as described in the method, with the following L-arginine concentrations: 0, 1 and 2 mM (lane 1-2) (**D**) Intensities of gel bands were quantified and plotted against the concentration of L-arginine. (**E**) Cartoon representation of TkHSP20 generated by AlphaFold2 [44]. (**F**) Plots of chemical shift perturbations of ‘hot spots’ of 50 μM [^13^C]-labeled *Pfu* pol upon adding 50 μM TkHSP20. (**G**) PCR amplification of 10 kb plasmid DNA by the *Pfu* pol in the presence of different concentrations of TkHSP20 from 0.1 pg/μl to 250 pg/μl. The PCR assays were performed as described in the methods, with the following TkHSP20 concentrations: 0, 0.1, 0.3, 1, 3, 10, 20, 50, 75, 100, and 250 pg/μl (lane 1-11) (**H**) Intensities of gel bands were quantified and plotted against the concentration of TkHSP20. Results expressed as mean ± SD of 3 independent biological replicates. Data were analyzed using unpaired Student’s test, ^**^P < 0.01.

To further investigate the application of chaperones in PCR, we cloned and expressed the small heat shock protein HSP20 from *Thermococcus kodakaraensis* (TkHSP20), which is conserved across species and functions in an ATP-independent manner (Fig. 2E) [32–35]. CSP analysis revealed that TkHSP20 interacts with *Pfu* pol at residues aro2, aro3, and aro4, covering three of the five active sites. Additionally, a significant decrease in the intensity of ali1 suggested its involvement in intermediate exchange interactions with TkHSP20 (Fig. 2F and S1B). Similar to L-arginine, TkHSP20 did not enhance the amplification of short DNA fragments (Fig. S2B). However, it significantly improved the amplification of long DNA fragments in a dose-dependent manner, with the most pronounced effect at 10 pg/μl, increasing the final PCR yield by ∼4-fold (Fig. 2G and H).

### 3.3 The mechanism of L-arginine and TkHSP20 in long-range PCR

The processivity of a DNA polymerase is a key determinant of its synthesis rate and substrate affinity, defined as the number of nucleotides incorporated during a single binding event. It can be indirectly assessed by measuring the amount of newly synthesized double-stranded DNA over a given period [39]. To evaluate the impact of L-arginine on *Pfu* pol processivity, we monitored nucleotide incorporation using EvaGreen fluorescence in a hairpin DNA template throughout the PCR reaction. The addition of L-arginine resulted in a decrease in the Michaelis constant (*K*_*m*_) from 0.87 ± 0.14 µM to 0.81 ± 0.08 µM, and revealed a slightly enhanced *k*_*cat*_ value from 128.86 ± 11.55 to 146.58 ± 7.68 s^−1^ (Fig. 3A). These findings imply that L-arginine moderately enhances the processivity of *Pfu* pol, which may contribute to its previously observed positive effect on PCR amplification efficiency.

**Figure 3.**
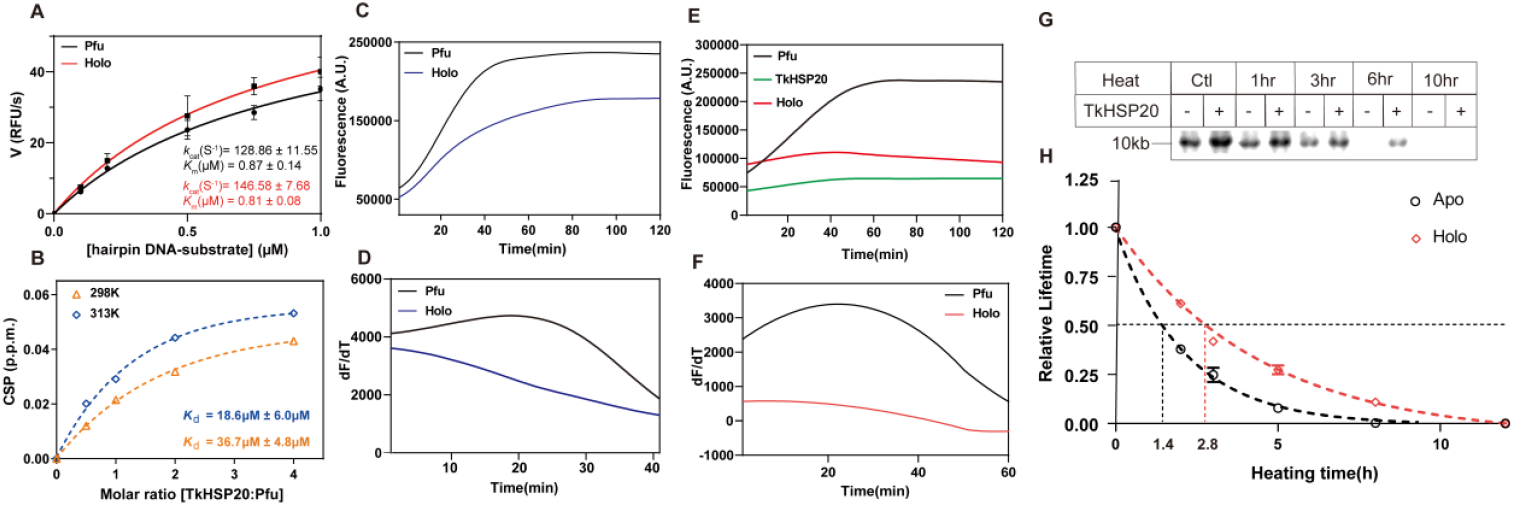
(**A**) Initial reaction rates (V_0_) of the *Pfu* pol were plotted versus different concentrations of the hairpin DNA substrate in the absence (black) and presence (red) of 2 mM L-arginine. Assays contained 2 nM of *Pfu* pol and different concentrations (0.1, 0.2, 0.5, 1 and 2 μM) of hairpin DNA templates respectively. The error bars represent the standard error of the mean from two biological replicates with three technical replicates for each biological replicate. The data was fitted to the Michaelis–Menten kinetics model. TkHSP20 improves the thermostability of the *Pfu* pol. (**B**) The magnitude of normalized CSP at 298K (orange) and 313K (blue) for the residue aro3 of 50 μM *Pfu* pol was plotted as a function of the molar concentration ratio of TkHSP20 and *Pfu* pol. Binding *K*_d_ was fitted for aro3. Mean *K*_d_ ± S.D. was 36.7 ± 4.8 μM (298K) or 18.6 ± 6.0 μM (313K). (**C** and **D**) Protein thermal shift (PTS) assay of *Pfu* pol in the presence (blue) and absence (black) of 20 mM L-arginine. The commercial dye SYPRO Orange was used to quantify the thermodynamic stability of *Pfu* pol during the continuous heating at 98 °C for 120 mins. The first derivative of the protein thermos shift plotted in **(D)**. (**E** and **F**) Protein thermal shift (PTS) assay of *Pfu* pol in the presence (red) and absence (black) of 30μM TkHSP20. The commercial dye SYPRO Orange was used to monitor the thermodynamic stability of the *Pfu* pol during the continuous heating at 98 °C for 120 mins. The first derivative of the SYPRO Orange fluorescence is plotted in panel (**F**). (**G)** PCR amplification of 10 kb DNA fragment, by using the *Pfu* pol after heating at 72 °C for 0, 1, 3, 6 and 10 hours (lane 1-2, lane 3-4, lane 5-6, lane 7-8, lane 9-10) in the presence and absence of 100 pg/μl TkHSP20 respectively. (**H**) Intensities of gel bands were quantified with Image J and plotted against the time of *Pfu* pol heated at 72 °C. The lifetime of *Pfu* pol is represented by its activity in amplifying 10 kb DNA fragments. Results expressed as mean ± SD of 3 independent biological replicates.

To further investigate the mechanism by which TkHSP20 facilitates long-fragment PCR with *Pfu* pol, we quantified the interaction between TkHSP20 and *Pfu* pol at different temperatures. NMR titration experiments were conducted by titrating TkHSP20 (0–200 µM) into 50 µM *Pfu* pol at 298K and 313K using a 600 MHz spectrometer (Fig. S3). The chemical shift perturbations (CSPs) of residue aro3 were fitted to determine the apparent dissociation constant (*K*_d_), revealing a stronger interaction at 313K (*K*_d_ = 18.6 ± 6.0 µM) than at 298K (*K*_d_ = 36.7 ± 4.8 µM). This suggests that TkHSP20 binds more tightly to *Pfu* pol at elevated temperatures, potentially preventing heat-induced denaturation (Fig. 3B). To confirm this, a protein thermal shift (PTS) assay was performed by continuously heating *Pfu* pol at 98°C in the presence and absence of TkHSP20 and L-Arginine respectively, with SYPRO Orange fluorescence used to monitor protein denaturation. The presence of both TkHSP20 and L-Arginine conferred significant thermal protection, as indicated by the minimal increase in fluorescence over 120 minutes, whereas *Pfu* pol alone exhibited substantial denaturation, with fluorescence intensity increasing approximately two-fold (Fig. 3C-F). The first derivative analysis of the PTS data further confirmed that TkHSP20 delayed *Pfu* pol denaturation (Fig. 3E and F). Additionally, an accelerated aging test at 72°C demonstrated that TkHSP20 extended the half -life of *Pfu* pol activity in amplifying a 10 kb fragment from 1.4 hours to 2.8 hours (Fig. 3G and H).

### 3.4 L-arginine and TkHSP20 exhibit a synergistic enhancement in long-range DNA amplification

The above results highlight the distinct yet potentially complementary roles of L-arginine and TkHSP20 in enhancing the performance of *Pfu* pol. To prove this, PCR reactions with a template of 10 kb were performed with L-arginine and TkHSP20. The results revealed that both L-arginine and TKHSP20 independently enhance the long-fragment amplification capability of *Pfu* Pol, while their combined application demonstrated a more pronounced enhancement in the amplification of 10 kb DNA fragments (Fig. 4A and B). This suggests that these two additives cooperatively improve *Pfu* Pol performance through a synergistic mechanism. For future applications in long-fragment DNA amplification, concurrent employment of arginine and TKHSP20 could be implemented to optimize reaction efficiency.

**Figure 4.**
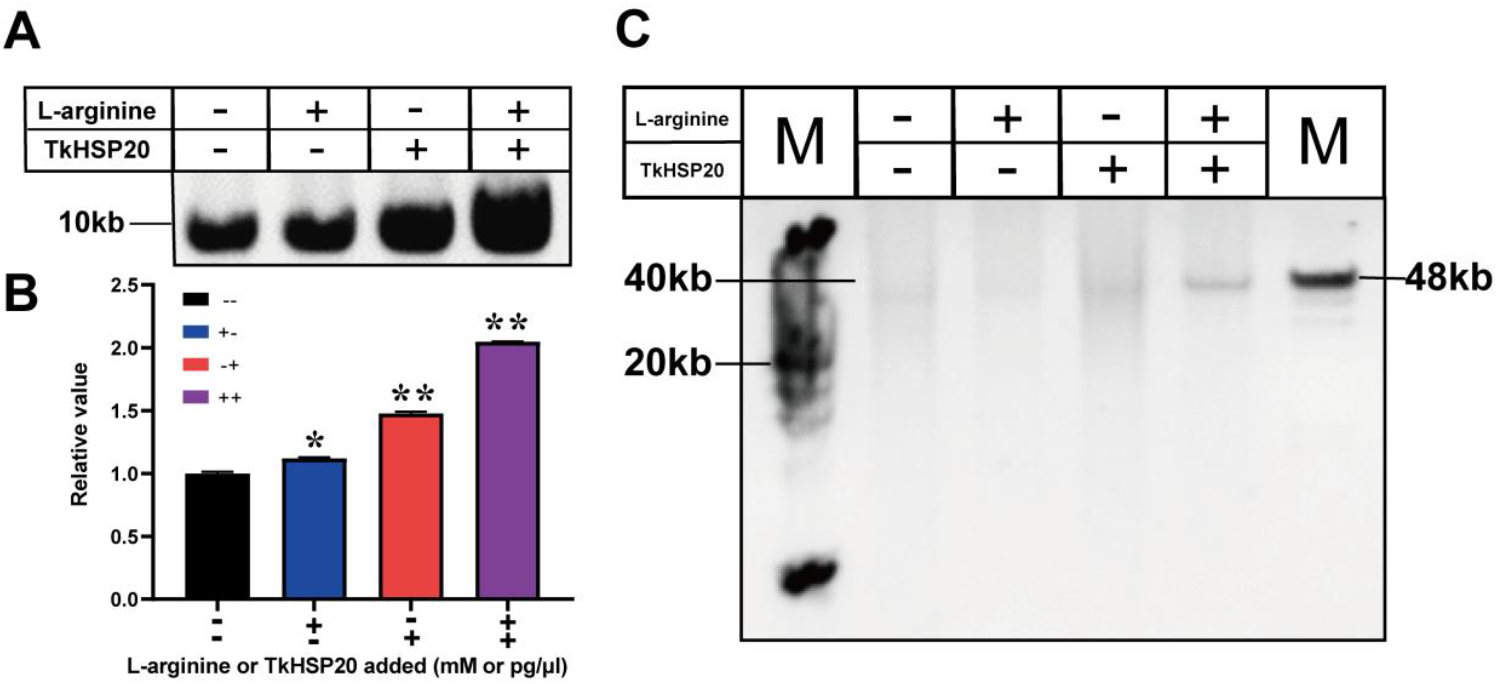
PCR amplification of 10 kb plasmid DNA (**A**) (**B**) or 40 kb λ DNA fragment (**C**) by the *Pfu* pol-S pol in the presence of 2 mM L-arginine or 3 ng/μl TkHSP20 separately or combinedly. The PCR assays were performed as described in the methods. Quantification of the intensity of gel bands was done with Image J. The intensity of the PCR product was plotted accordingly. Results expressed as mean ± SD of 3 independent biological replicates. Data were analyzed using unpaired Student’s test, ^*^ P< 0.05, ^**^P < 0.01.

## 4. DISCUSSION AND CONCLUSION

Although plenty of PCR-derived techniques have been devised and widely adopted across various domains of molecular biology and biotechnology. The limitations related to long-range amplification inherent in current PCR methodologies impede a broader utilization of these related techniques. The challenges in amplifying lengthy DNA fragments involve several factors, which may intensify with more complex templates [36]. Firstly, prolonged heating during PCR can compromise the activity of DNA polymerases, which must remain catalytically active throughout the PCR process to support DNA amplification. Secondly, high temperatures can also lead to the degradation of DNA or newly synthesized amplicons to the point where templates become severely fragmented for amplification [37]. Thirdly, as the length of DNA increases, the likelihood of forming secondary structures and premature amplicon increases which may further hinder PCR [38]. Finally, long-range PCR is sensitive to even minor variations in reaction conditions, which may not always be obvious and, thus, are challenging to detect. Developing a long PCR-based assay can be a burdensome and unrewarding endeavour [39]. Here, we employed NMR spectroscopy to identify both TkHSP20 and L-arginine could improve the activity of the *Pfu* pol in amplifying long DNA fragments. Especially, TkHSP20 almost completely prevents the denaturation of the *Pfu* pol at 98 °C for 2 hours, significantly enhancing thermostability and processivity of *Pfu* pol. The chemical L-arginine, can not only stabilize proteins but also nucleic acid [40]. To overcome the problem of unwanted secondary structure formation and premature amplicon, a recent suppression thermo-interlaced (STI) PCR method was developed to repress the amplification of smaller non-specific products [41]. Using the STI-PCR method, we observed clear amplified PCR bands of 40 kb DNA fragments with the presence of TkHSP20 and L-arginine, while the control group failed by using solely *Pfu* pol (Fig. 4C). Additionally, the increased stability of the DNA polymerase by TkHSP20 also showed an obvious potential in the application for storage and handling of DNA polymerase, which may also be extended to use as an additive for preserving and transporting other heat-sensitive enzymes.

The NMR-based approach for discovering new additives or co-factors for DNA polymerase in our work is innovative in its clarity and generalizability. This versatile method can be extended to optimize other enzymes, including co-factor identification and buffer optimization. It doesn’t rely on the pre-determined assignment information. Measurements of ^1^H-^13^C moieties extended the size of proteins suitable for NMR spectroscopy up to Mega-Dalton [42], enabling the identification of active sites of high-molecular-weight enzymes. Compared to backbone amide signals, side chain signals exhibit greater sensitivity in detecting subtle chemical environmental changes induced by weak molecular interactions, consequently manifesting more pronounced chemical shift perturbations. This enhanced sensitivity facilitates the identification of cofactors with weak interactions, such as chaperones. This NMR guided approach provides the experimentally determined hots spot and it is not dependent on the structural data or bioinformatics, compared to the current computational methods for predicting hot spots. Furthermore, these determined hot spots could not only be used as probes for rational optimization of the target enzyme involved reactions but also enable the full potential of directed enzyme evolution with the assignment information [25,43]. Last but not least, the newly identified TkHSP20 and L-arginine in our work facilitate the application of *Pfu* pol in the lengthy DNA fragment PCR, benefiting various cloning, functional analysis, and nanopore based long fragment sequencing and representing a significant technological advancement that promises to accelerate the fields of synthetic biology, molecular biology, and biotechnology.

## CRediT authorship contribution statement

Y.C., M.Z., R.D., and L.H. planned the study and designed experiments. Y.C., M.Z., and R.D. performed experiments. X.Z., M.L., and L.H. acquired funds. All authors analyzed and interpreted the results. Y.C., M.Z., R.D., and L.H. wrote the paper with input from all authors.

## Appendix A. Supplementary data

Supplementary material.

## Data Availability

Data will be made available on request.

