## Supplemental data for "NMR-guided rational exploration of co-factors in boosting the *Pfu* DNA polymerase"

**SUPPLEMENTARY DATA**

**Table S1.** Synthesized oligonucleotides.

| | |
|---|---|
| Hairpin DNA | 5’-CCAGCATTATGAAAGTGACACGTGCACCATTGGTGCACGTG-3’ |
| 10K F | 5’-ATCCTGGACGTGGATTACATCACC-3’ |
| 10K R | 5’-CATATGACGACCTTCGATATGGCC-3’ |
| Ap cDNA F | 5’-GTCCTGGCTCTCTTCTACAAGG-3’ |
| Ap cDNA R | 5’-GGTACGTGCTGTTGAACTGTTCC-3’ |
| Mt gDNA F | 5’- AACCCAGAAGACAGGTGGAAAG-3’ |
| Mt gDNA R | 5’- GCCAGATTACGTATATCCTGGCAG-3’ |
| M13 F | 5’-AAGCCATCCGCAAAAATGACCTCT-3’ |
| M13 R | 5’-GTCAGAAGCAAAGCGGATTGCATC-3’ |

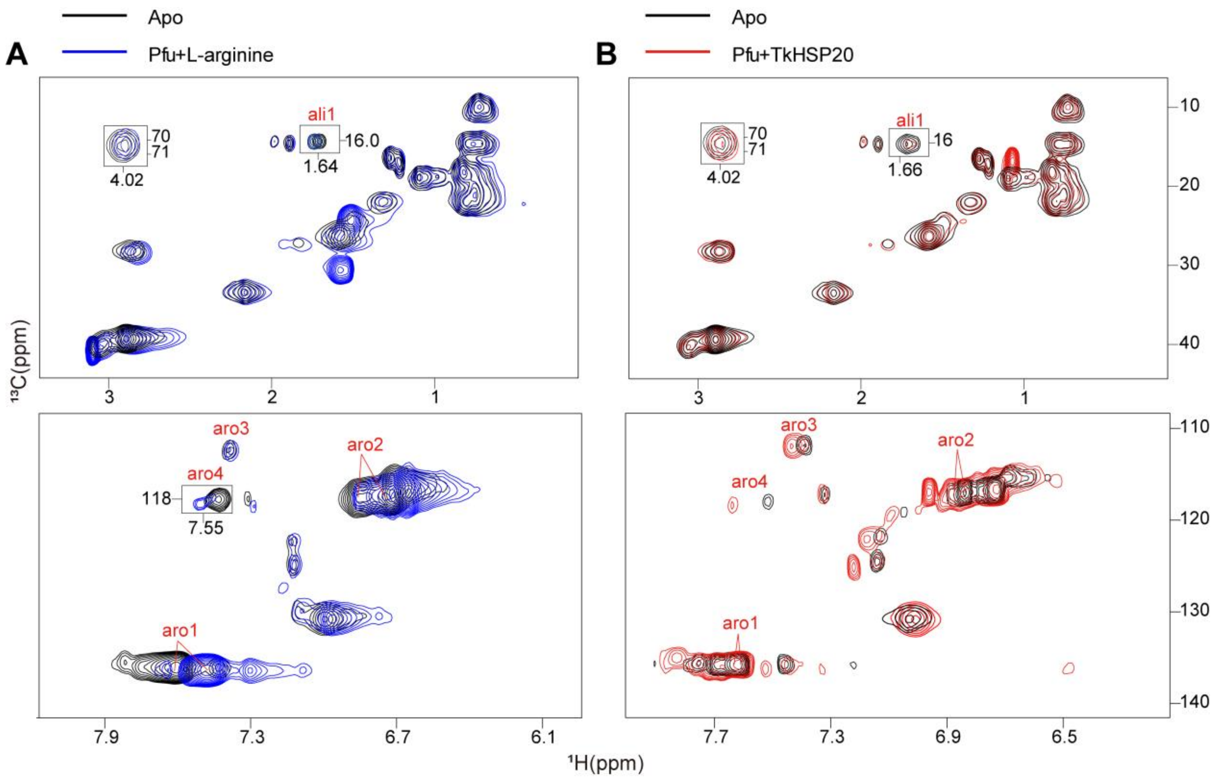


**Figure S1.** 2D [$^{13}$C,$^{1}$H]-HMQC spectra of 50 μM [$^{13}$C]-labeled *Pfu* pol acquired on the 600 MHz ($^{1}$H frequency) spectrometer at 298 K in the absence and presence of 20 mM L-arginine (**A**) or 50 μM TkHSP20 (**B**).

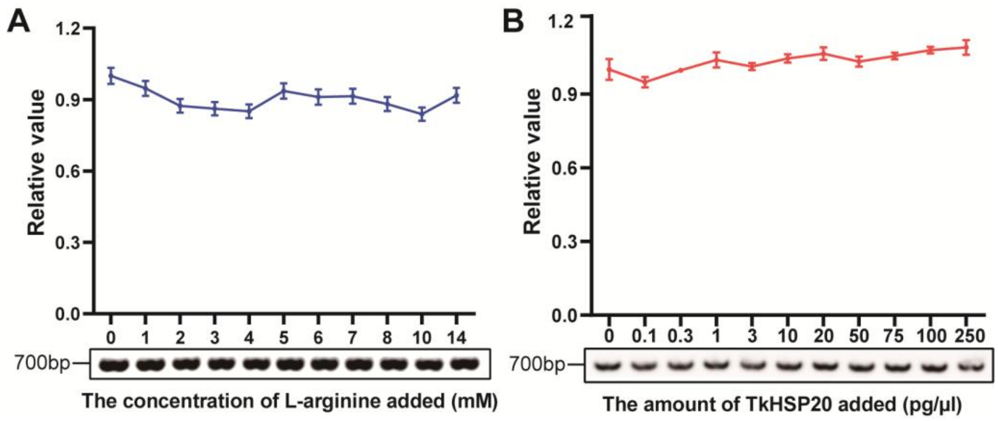


**Figure S2.** PCR amplification of mouse tail gDNA extracts (with target length 700 bp) by the *Pfu* pol polymerase in the presence of different concentrations of L-arginine (**A**) from 1 mM to 14 mM or the different concentration of TkHSP20 from 0.1 to 250 pg/μl (**B**). The PCR assays were performed as described in the methods. Intensities of gel bands were quantified and plotted against the concentration of L-arginine or ThHSP20, showing on top of the corresponding gel image. Results expressed as mean ± SD of 3 independent biological replicates. Data were analyzed using unpaired Student's test, **P < 0.01.

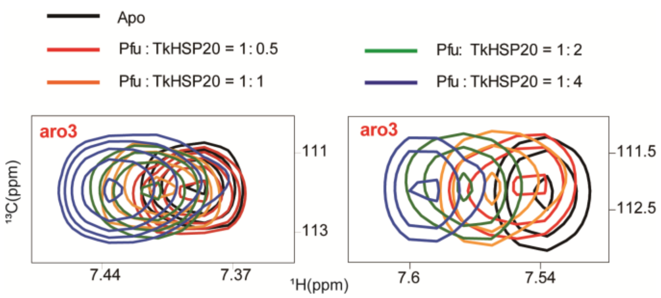


**Figure S3.** Peak of aro3 of the *Pfu* pol polymerase are showed separately to reveal the chemical shift perturbations (298K, Left, 313K, Right) upon adding different concentrations of TkHSP20 (0 μM black, 25 μM red, 50 μM orange, 100 μM green, 200 μM blue) respectively.
